# Single-agent Foxo1 inhibition normalizes glycemia and induces gut β-like cells in streptozotocin-diabetic mice

**DOI:** 10.1101/2022.03.26.485929

**Authors:** Yun-Kyoung Lee, Yaohui Nie, Bryan Diaz, Nishat Sultana, Takumi Kitamoto, Wen Du, Rudolph L. Leibel, Domenico Accili, Sandro Belvedere

## Abstract

Insulin treatment remains the sole effective intervention for Type 1 Diabetes. Here, we investigated the therapeutic potential of converting intestinal epithelial cells to insulin-producing, glucose-responsive β-like cells by targeted inhibition of Foxo1. We have shown that this can be achieved by genetic ablation in gut Neurogenin3 progenitors, adenoviral or shRNA-mediated inhibition in human gut organoids, and chemical inhibition in Akita mice, a model of insulin-deficient diabetes. In the present study, we provide evidence that two novel Foxo1 inhibitors, FBT432 and FBT374 have glucose-lowering and gut β-like cell-inducing properties in mice rendered insulin-deficient by administration of streptozotocin. FBT432 is also highly effective in combination with a Notch inhibitor in this model. The data add to a growing body of evidence suggesting that Foxo1 inhibition be pursued as an alternative treatment to insulin administration in diabetes.

## Introduction

*Type 1* diabetes (T1D) is a partially unmet medical need, with an estimated 18 million patients worldwide requiring life-long insulin supplementation, as do many patients with *type 2* diabetes (T2D) due to loss of adequate endogenous insulin secretion (1). Since the introduction of insulin treatment in 1922, there have been improvements in composition, formulation, dosing regimens and devices. Nevertheless, the human burden of treatment remains high, with over half of the patients failing to achieve optimal glucose control and experiencing complications (2; 3).

There has been significant progress in delivering insulin-producing cells generated by differentiation of either embryonic stem or induced pluripotent stem cells to T1D patients (4). This approach has the potential to restore autonomous glucose control and minimize the side effects of insulin administration (e.g. hypoglycemia, weight gain) (5; 6), but there is lingering uncertainty on the mechanics of delivering a sufficient number of exogenous cells to patients and protecting them from the attendant immune responses.

An attractive alternative is in vivo generation of insulin-producing, glucose-responsive β-like cells from other cell types. We and others have demonstrated the feasibility of this approach by reprogramming enteroendocrine cells (EECs) (7). Inhibition of Foxo1 by genetic ablation induced the formation of β-like cells *in vivo* in streptozotocin-diabetic mice, restoring glucose control (8), and inhibition by shRNA or a dominant negative mutation induced β-like cell conversion in human iPS-derived gut organoids *in vitro* (9).

More recently, we have demonstrated that this new therapeutic paradigm can be realized by oral treatment with a small-molecule tool compound Foxo1 inhibitor (FBT10), when used in combination with a Notch inhibitor to increase the pool of EECs available for conversion to insulin-producing cells. We observed a significant improvement in glucose levels and glucose tolerance in Akita mice, a model of insulin-deficient diabetes (10).

Here we report that two novel, orally bioavailable, optimized Foxo1 inhibitors, FBT432 and FBT374, can lead to the formation of gut β-like cells, normalize glucose levels and glucose tolerance in STZ diabetic mice either as single agents, or in combination with a Notch inhibitor. Specifically, compound FBT374, when administered as a single agent, normalized blood glucose concentrations and restored a wildtype-like glucose tolerance pattern. Taken together, these results suggest that Foxo1 inhibition is a viable strategy for T1D treatment.

## Research Design and Methods

### Chemicals

FBT374 and FBT432 were synthesized by IntelliSyn Pharma, (Montreal, Canada) and determined to be > 98% pure by LC/MS and ^1^H NMR spectroscopy. PF-03084014 was synthesized by Viva biotech (Shanghai, China) according to published methods.

### *In vitro* ADME and *in vivo* pharmacokinetic and tissue distribution

Microsomal Stability was determined by incubating compounds at 0.3 μM in 0.1 M PBS (pH 7.4, 0.015 % DMSO) in duplicate at 37°C, in the presence of 0.25 mg/mL mouse liver microsomes (BioIVT) and 2 mM NADPH. Aliquots of the reaction were quenched at 0, 5, 15, 30, 45 minutes, and analyzed by LC-MS/MS (Agilent 1290 tandem with Sciex 5500 Qtrap). Kinetic solubility was determined in duplicate by diluting a DMSO stock of test compounds into aqueous phosphate buffer (pH 7.4, 1% final DMSO) at a concentration of 100 μM. The solution was shaken for 1h, filtered and analyzed by LC-MS/MS (Agilent 1290 tandem with Sciex 5500 Qtrap) against a calibration curve. Cell permeability was measured by adding the test compound (2 μM in HBSS and 0.4% DMSO) either to the apical or basolateral side of canine MDR1 Knockout MDCKII Cells for 90 minutes at 37°C. Apical and basolateral solutions were analyzed by LC-MS/MS (Agilent 1290 tandem with Sciex 5500 Qtrap). *In vivo* PK was performed in male ICR (n=3 per route). FBT374 or FBT432 were formulated in Solutol HS-15: Saline = 5:95 (v/v). Mice were dosed i.v. at 1 mg/kg or p.o. at 10 and 50 mg/kg, and plasma isolated from blood collected at 0.083, 0.25, 0.5, 1, 2, 4, 8, and 24-hr. Tissues were collected 1 and 8 hr after heart perfusion with 1x PBS for 1 min and homogenized in PBS (1:3, w/v). Samples were analyzed by LC-MS/MS (Agilent 1290 tandem with Sciex 5500 Qtrap).

### Reporter gene assays

Reporter gene assays were performed as previously (11). Briefly, HEK293 cells were seeded at 7,500 cells/well in EMEM supplemented with 1% fetal bovine serum (FBS) and 1x penicillinstreptomycin (Gibco) onto 384-well plates. After overnight incubation, cells were transfected with pGL.4.26-4xIRE-luc2 (Fluc), pRL-CMV (Promega), and pcDNA3.1 vector containing RFP (as negative control), FOXO1-AAA (Addgene, #9023), FOXA2 (GenScript, #OHu31644), FOXO3 (Genscript, #OHu23372), FOXO4 (GeneScript, #OHu23105) using Lipofectamine3000 (Invitrogen). Cells were treated with 10-point-half-log dilution of compound in duplicate wells at a final DMSO concentration of 0.5%. After 24hr, lysates were used to determine reporter gene activities using Dual-Glo Luciferase Assay (Promega) and medium from the plate was used to measure cytotoxicity using LDH-Glo Assay (Promega) with an EnVision 2105 plate reader (Perkin Elmer). At least two independent experiments were performed for each compound for each assay. IC_50_ or LC_50_ was estimated by the four-parameter logarithmic curve fit using GraphPad Prism.

### Primary hepatocyte isolation and qPCR

All procedures were approved by the CUNY-Advanced Science Research Center (ASRC) Institutional Animal Care and Use Committee (IACUC). Hepatocytes were isolated from 8- to 10-week-old male C57/BL6 mice as described (11). The isolated hepatocytes were seeded at 150,000 cells/well in M199 medium (Gibco) containing 10% FBS, 1x penicillin-streptomycin, and 50 mg/ml of G418 (Gibco) onto collagen-coated 24 well plates. After 15-16h in M199 medium supplemented with 1% FBS, cells were treated with freshly prepared 100 mM cAMP (Sigma), 1 mM dexamethasone (Sigma) and concentrations of FBT ranging from 30 mM to 100 nM or DMSO for 6 hr. RNA was extracted from using QIAsymphony (Qiagen) and reverse-transcribed using 0.1mg and the High-Capacity cDNA Reverse Transcription Kit (Applied Biosystem). qPCR was performed on a CFX Connect Real-Time PCR system (Bio-Rad) using iTaq Universal SYBR Green SuperMix. Primer sequences are indicated in Supplementary Table 4.

### Pyruvate tolerance test (PTT)

6- to 7-week-old C57/BL6 male mice (Jackson Laboratories) were acclimated for 1 week. FBT374 or FBT432 was formulated into *N,N*-Dimethylacetamide: Solutol HS 15: water=5:10:85 (v/v/v) solution, pH 4-5. Mice were randomized based on weight and dosed at 15 and 50 mg/kg/dose, 10 ml/kg b.i.d. p.o. for a total of 3 doses (The third and last dose was given 2 hours prior to pyruvate injection). After a 4-hr fast, 2g/kg of sodium pyruvate (Sigma) dissolved in saline was injected i.p. at 10 ml/kg and the glucose levels were measured at 0, 15, 30, 60 and 120 min by tail vein sampling using Contour Next glucometer (Bayer).

### Animal study using streptozotocin (STZ) induced diabetic mice

6- to 7-week-old C57/BL6 male mice were purchased from Jackson Laboratories and acclimated for a week, and injected i.p. with STZ dissolved in sodium citrated buffer (pH4.5) after 4- to 5-hr fast. Glucose was monitored daily until mice became hyperglycemic. Mice were implanted subcutaneously with a half pellet of LinBit (LinShin Canada). Once hyperglycemia stabilized ~600mg/dl, animals were randomized based on weight and ambient glucose. FBT374 or FBT432 was formulated into *N,N*-Dimethylacetamide: Solutol HS 15: water= 5:10:85 (v/v/v) solution, pH4-5 and dosed orally twice daily at 50 mg/kg/dose, 10 ml/kg. For combined treatment with FBT432 and PF-03084014, the solution was prepared in *N,N*-Dimethylacetamide: Solutol HS 15: water= 5:10:85 (v/v/v) solution, pH 4-5 to achieve dosing concentration of 150 and 50 mg/kg for PF-03084104 and FBT432, respectively. Weight was monitored daily and circulating glucose concentration in a blood drop obtained from a small nick on the lateral tail vein using a 30G needle was monitored using a glucometer (Nova Medical) every 2-3 days in awake mice. For glucose tolerance tests, mice were fasted for 4-hr (8:30 am – 12:30 pm) and dosed i.p. with 2 g/kg of D-glucose (Sigma) dissolved in saline. Glucose was measured at the indicated time points as indicated above. Urine ketone and glucose were measured using Diastix (Bayer). Mice were killed by CO2 inhalation followed by cervical dislocation. Of note, we observed an increased size and volume of the cecum following FBT374, FBT432, or combination treatment. The tissue was intact and healthy with no sign of increased of hyperplasia, hypertrophy, or inflammation (Supplementary Figure 1). Small and large intestines were collected for immunohistochemistry and epithelial cell isolation followed by flow cytometry and qPCR. Liver paraffin sections were stained with hematoxylin and eosin (H&E). Images were taken using the Keyence BZ-X800 microscope. RNA was extracted from snap-frozen liver using QIAsymphony (Qiagen) and reverse-transcribed using 0.5mg and the High-Capacity cDNA Reverse Transcription Kit (Applied Biosystem). qPCR was performed on a CFX Connect Real-Time PCR system (Bio-Rad) using iTaq Universal SYBR Green SuperMix. Primer sequences are indicated in Supplementary Table 4.

### Epithelial cell isolation and EEC analysis

The proximal third of the small intestine was used for epithelial cell isolation as described (10). Briefly, the cut-open and washed intestinal tissue segment was incubated in 5 ml of 30 mM EDTA/1.5 mM DTT/DPBS on ice for 20 min, followed by incubation in 5 ml of 30 mM EDTA/DPBS at 37°C for 8 min. After vigorous shaking for 30 sec, pellets containing villi and crypts were collected by centrifugation at 600xg for 10 min. Pellets were resuspended in HBSS containing 10 mM of Rock inhibitor (Y-27632, SelleckChem) and 0.3 U/ml of Dispase (Roche) and incubated at 37°C for 7 min with vigorous shaking at 5 and 7 min. The cell suspensions were sequentially passed through 70 mm and 40 mm filters (Fisher) and the single cell suspension was collected. Cells were then lysed in RLT buffer for RNA isolation using QIAsymphony or immunostained with ChgA, 5HT, and GLP-1 antibodies followed by flow cytometry on a BD LSR FortessaTM Cell Analyzer as previously.

### Immunostaining and quantification

The small and large intestines were dissected, rinsed in 1x PBS and fixed with 4% paraformaldehyde (PFA)/PBS by passing through the solutions. Tissues were cut open, washed, fixed in 20 ml of 4% PFA/PBS for 90 min at 4°C on a rocker, dehydrated in 30% sucrose/PBS for 24 hr, then fashioned into swiss rolls of small and large intestines, embedded in OCT (Sakura), and cut into 6-7 μm-thick frozen sections. The list of primary and secondary antibodies is shown in Supplementary Table 5. Images were taken using the Keyence BZ-X800 microscope and quantification of 5HT- and insulin-immunoreactive cells was performed using Hybrid Cell Count software (Keyence).

## Results

### FBT432 activity in cell-based assays and in vivo efficacy

Based upon the previously reported Foxo1 inhibitor FBT10 (10; 12), we generated a series of chemical variants to optimize its properties and identified FBT432, a compound with improved solubility and *in vitro* metabolic stability (Supplementary Table1). We evaluated activity and selectivity of FBT432 against FOXO in previously established transcription reporter gene assays by transiently expressing FOXO1 and an insulin response element (IRE)-firefly luciferase construct as a reporter of FOXO1 activity in HEK293 cells. We used FOXA2 as a counter screen to determine the compound’s selectivity as FOXA2 binds to the same DNA elements but is not responsive to insulin signaling (13). As shown in Table 1, FBT432 showed strong activity against FOXO1 (estimated IC_50_= 69 nM) but no activity against FOXA2 and other FOXO isoforms, including FOXO3 and FOXO4. A further test using a CMV-promoter mediated Renilla luciferase (Rluc) reporter gene showed no activity, revealing FBT432 as a potent and specific FOXO1 inhibitor. FBT432 showed no significant cellular toxicities (Table 1).

**Table 1.** *In vitro* activities of FBT432 and FBT374 in HEK293 cell-based assays. IRE: insulin response element. Rluc: *Renilla* luciferase. Data are geometric mean x /÷ standard deviation.

|  |  | FBT432,<br>IC<sub>50</sub> (μM) | FBT374,<br>IC<sub>50</sub> (μM) |
| --- | --- | --- | --- |
| IRE-reporter assays | FOXO1 | 0.069 ×/÷ 1.79 | 0.073 ×/÷ 1.20 |
|  | FOXO3 | >50 | 16.2 ×/÷ 1.01 |
|  | FOXO4 | >50 | 1.63E+01 |
|  | FOXA2 | >50 | >50 |
| Non-specific activities | pCMV-Rluc | >50 | >50 |
|  | Cell viability (LC<sub>50</sub>) | >50 | > 50 |

Next, we examined the effect of FBT432 on endogenous FOXO1 function using mouse primary hepatocytes as previously described (11). Five-point dose-response treatment of primary hepatocytes with FBT432 showed an inhibition of cyclic AMP (cAMP) and dexamethasone (Dex)-induced *G6pc* expression at concentrations above 1 μM, with an estimated IC_50_ of 5.28 μM (Figure 1A).

**Figure 1.**
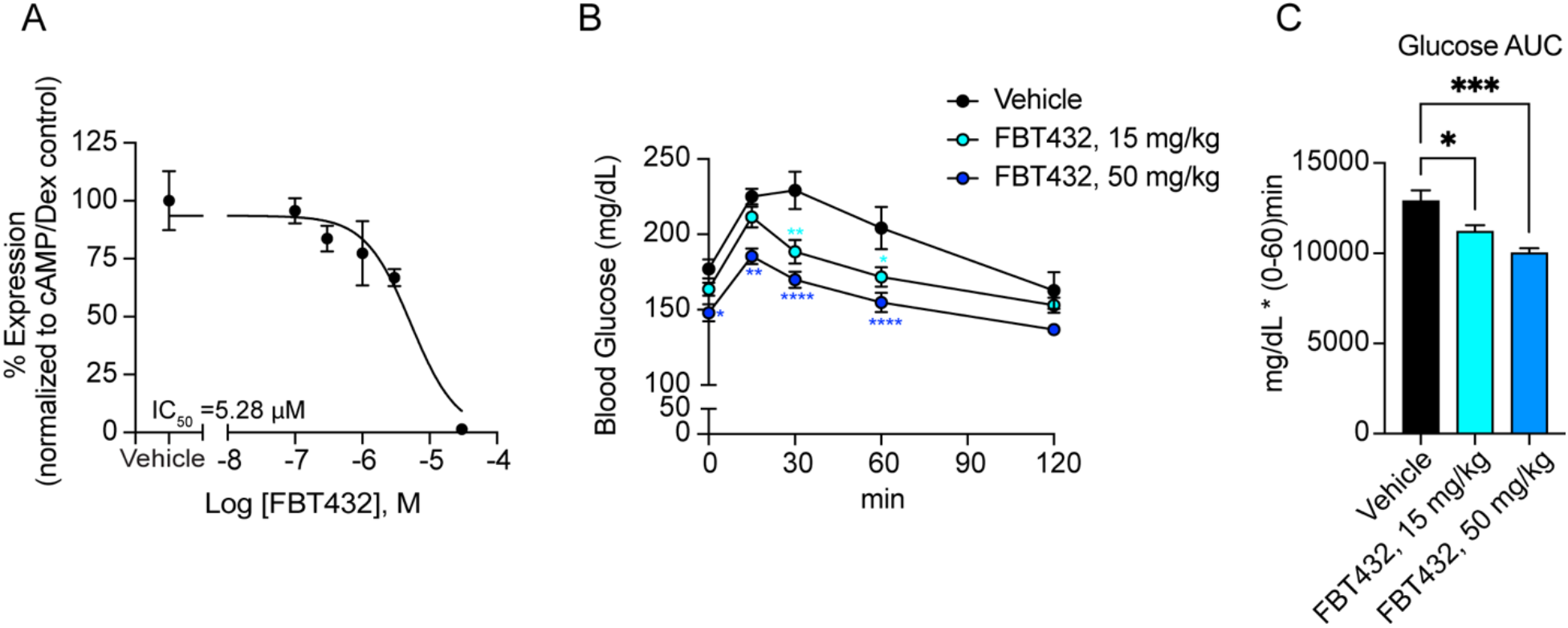
*In vitro* and *in vivo* activity of FBT432. (A) *G6pc* mRNA in hepatocytes treated with the indicated combinations of reagents. Data are means ± SEM of 2-3 replicate wells per concentration. IC_50_ was generated by four-parameter curve fit in GraphPad Prism. (B, C) Glucose levels and area-under-the-curve (AUC) during PTT after treatment with FBT432. N= 8 mice per group; *, **, ***, ****=p < 0.05, 0.01, 0.001, 0.0001 vs. vehicle by two-way ANOVA (B) or oneway ANOVA (C).

FBT432 is orally bioavailable in mouse, with good tissue distribution, including liver and intestine (Supplementary Table 2). To evaluate its *in vivo* efficacy, we used pyruvate tolerance tests (PTT) as a pharmacodynamic (PD) measure of the compound’s ability to suppress hepatic glucose production, an acute consequence of FOXO1 inhibition (11). Normoglycemic mice received three oral doses at 15 and 50 mg/kg on a b.i.d. schedule, and glucose levels were monitored at 0, 15, 30, 60, or 120 min after a bolus intraperitoneal injection of pyruvate following a 4-hour fast. Mice treated with FBT432 showed a dose-dependent lowering of glucose excursions compared to vehicle-treated mice (Figure 1B-C). These results suggest that FBT432 is a bioavailable, potent, and selective FOXO1 inhibitor, leading us to further evaluate its *in vivo* efficacy in a preclinical model of diabetes.

### Glucose-lowering by FBT432 in streptozotocin (STZ) diabetes

We have shown that somatic ablation of Foxo1 in Neurogenin3 (Neurog3)-expressing endocrine progenitor cells normalizes hyperglycemia by converting gut enteroendocrine cells (EECs) into β-like insulin-producing cells. We therefore tested whether chemical FOXO1 inhibition by FBT432 can lower glycemia in diabetic mice by converting EECs into insulin-producing cells. FBT10 lowers glucose in Akita mice when administered at 50 mg/kg on a b.i.d. schedule for five days (10). To rule out a confounding effect of residual insulin secretion as well as of the specific insulin gene mutation of Akita mice (14), we used streptozotocin, administered to 7- to 8-week-old C57/BL6 mice at 170 mg/kg intraperitoneally, to induce hyperglycemia, accompanied by glucosuria, ketonuria, near-absent plasma insulin and immunoreactive pancreatic insulin (data not shown).

To prevent diabetic ketoacidosis (DKA)-associated morbidity and mortality, we implanted a sub-therapeutic dose of insulin by way of a half pellet of LinBit. This expedient allowed us to maintain a stable hyperglycemia of ~600 mg/dl. After monitoring animals for up to 3 days, we administered FBT432 at 50 mg/kg/dose by gavage on a b.i.d. schedule for 12 days. Consistent with the lack of cellular toxicity observed *in vitro*, administration of FBT432 was well tolerated and caused no change in body weight (Figure 2A). FBT432 demonstrated a statistically significant glucose-lowering effect after 2 days of treatment that persisted until the end of the experiment on day 10 (Figure 2B). The fasting blood glucose concentrations of the FBT432-treated group were significantly lower than the untreated control group (Figure 2C). Moreover, intraperitoneal glucose tolerance tests (ipGTT) confirmed that FBT432 treatment had a significant glucose-lowering effect (Figure 2D, E). Accordingly, ketonuria was dramatically reduced, consistent with restored insulin action (Figure 2F).

**Figure 2.**
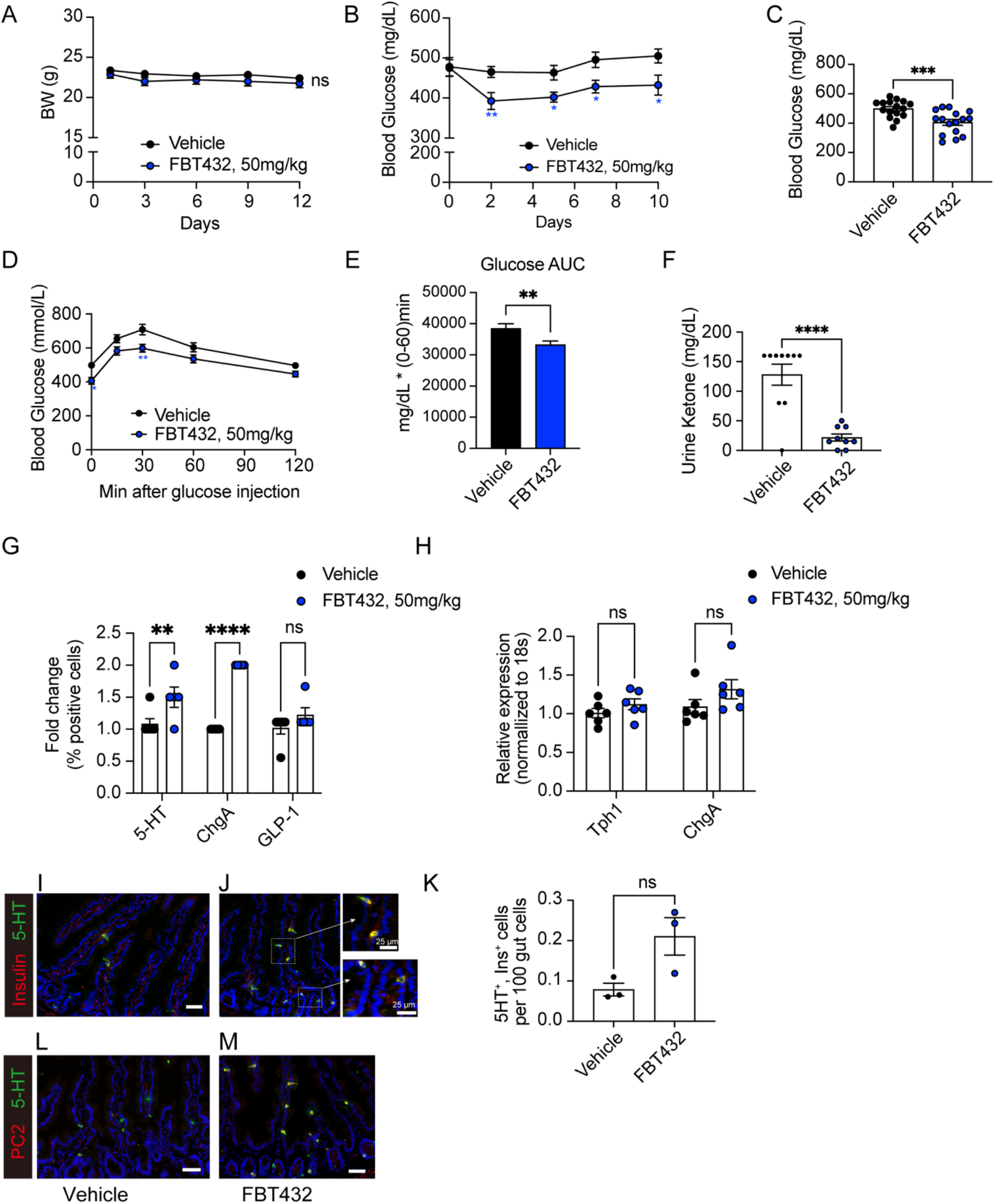
Effect of FBT432 in STZ diabetic mice. Mice were randomized as described and treated with vehicle or FBT432 (50 mg/kg/ b.i.d.) for 12 days. (A) Weight was measured before morning dosing. NS= not significant vs. vehicle by two-way ANOVA. N=9-16 per group. Aggregated data from 2 independent experiments conducted under the same condition and presented as means ± SEM. (B) *Ad libitum* glucose levels measured between 4 and 5 pm before afternoon dosing. *, **=p < 0.05, 0.01 vs. vehicle by two-way ANOVA. (C) 4-hr fasting glucose on day 12. ***= p < 0.001 by two-tailed t-test. (D) Glucose tolerance test (GTT) on day 12. *, **=p < 0.05, 0.01 vs. vehicle by two-way ANOVA. (E) AUC from 0-60 min during GTT. **= p < 0.01 vs. vehicle by two-tailed t-test. (F) Urine ketone (acetoacetic acid) concentration on day 13. ****= p < 0.0001, NS= not significant vs. vehicle by two-tailed t-test. (G) Flow cytometry of 5HT-, ChgA-, and GLP-1-immunoreactive cells from the proximal gut. N=3-4 per group, data are means ± SEM. **, ****= p < 0.01, 0.0001 vs. vehicle by two-way ANOVA. (H) qPCR in isolated epithelial cells. N=6 per group, data are means ± SEM. NS= not significant vs. vehicle by two-way ANOVA. (I-M) Representative intestinal immunohistochemistry at day 12 of treatment with vehicle or FBT432. (I, J) 5HT (green) and Insulin (red), (L, M) 5HT (green) and PC2 (red). Scale bar= 50 μm, DAPI counterstains nuclei. (K) Quantification of 5HT- and insulin-co-immunoreactive cells. N=3 per group, data are means ± SEM. NS= not significant vs. vehicle by two-tailed t-test.

To assess whether the glucose-lowering effect of FBT432 administration was driven by newly arisen gut insulin-producing cells, we analyzed the EEC pool. We isolated epithelial cells from the proximal gut, including duodenum and jejunum, immunostained them with Chromogranin A (ChgA), 5HT, and GLP-1 and analyzed each cell sub-population by flow cytometry. FBT432 treatment slightly but significantly increased total EEC number, as measured by counting ChgA cells. Interestingly, the 5HT-positive population was significantly increased, while the GLP-1-positive population remained unchanged (Figure 2G). Expression of marker genes of either sub-population was not significantly changed (Figure 2H).

Next, we asked whether EECs had been converted to β-like cells. We co-immunostained small intestine sections with insulin and 5HT antibodies. Consistent with the glucose-lowering effect, we observed the appearance of insulin- and 5HT-immunoreactive cells in FBT432-treated mice, mostly in duodenum (Figure 2I, J, K). Prohormone convertase-2 acts in pancreatic b-cells to process proinsulin to mature insulin and is not normally found in the intestine (8). To further characterize the insulin-positive gut cells, we tested PC2 immunoreactivity in the FBT432-treated group and found an increase that paralleled insulin-immunoreactive cells, mostly in duodenum. These PC2-positive cells were also co-immunoreactive with 5HT (Figure 2L, M).

### Combination treatment with Notch and Foxo1 inhibitors normalizes glycemia in STZ mice

Although FBT432 administration showed a glucose-lowering effect, it underperformed the genetic Foxo1 knockout. We have previously proposed that FOXO1 and Notch inhibitor combination therapy has synergistic effects, with Notch inhibition acting to promote EEC progenitor differentiation into the secretory cell lineage, thereby expanding the pool of cells available for conversion to β-like cells by FOXO1 inhibition (10). Consequently, we have shown that combination treatment with the first-generation Foxo1 inhibitor, FBT10, and the gamma-secretase inhibitor PF-03084014, had additive effects on β-like cell generation and systemic glucose homeostasis in Akita mice (10).

We further investigated whether it is possible to achieve a synergistic glucose lowering effect by combining Notch inhibition with the orally bioavailable clinical gamma secretase inhibitor PF-03084014 (PF) and FBT432 in the STZ model. The effects of treatment with PF alone are described in the previous report (10). Oral administration of PF at 150 mg/kg and FBT432 at 50 mg/kg b.i.d. for 10 days was well tolerated, with a slight decrease of weight (Figure 3A). A significant glucose-lowering effect occurred within 4 days of starting combination treatment, and led to a normalization of glycemia after 7 days of treatment (Figure 3B). We saw a similar effect on fasting glycemia (Figure 3C), and intraperitoneal GTT in mice treated with PF and FBT432 (Figure 3D, E). Urine glucose and ketone became undetectable, consistent with restoration of insulin action (Figure 3F, G). We analyzed the intestine of animals treated with the combination to assess the efficacy of FBT432 and PF in converting EECs into β-like cells. Insulin immunohistochemistry identified numerous cells, mostly localized to the base of crypts, and more evident in the proximal gut, (duodenum and jejunum; Figure 4A, B). We confirmed that insulin-positive cells co-expressed 5HT (Figure 4A, B), consistent with the result of single treatment with FBT432 (Figure 2K). Quantification of insulin-immunoreactive cells confirmed a significant increase by PF + FBT432 combination treatment compared to vehicle (Figure 4C). Other b-cell markers, such as PC2 and C-peptide, were also increased by combination treatment (Figure D-G). mRNA analyses in epithelial cells isolated from the proximal gut confirmed a significant induction of b-cell markers, such as *Ins2, Nkx6.1, Nkx2.2, Kir6.2*, and *Sur1* (Figure 4K). Consistent with expectations, combination treatment with PF-03084014 and FBT432 significantly enriched the 5HT EECs sub-population in the proximal gut, as assessed by immunostaining (Figure 4H, I) and flow cytometry (Figure 4J). Gene expression showed a significant increase of Neurog3 by combination treatment, consistent with an expanded secretory progenitor pool (Figure 4K). Collectively, these results are consistent with the hypothesis that FOXO1 inhibition drives EEC differentiation into β-like cells, and that this effect is significantly enhanced by Notch inhibition, leading to a substantial glucose-lowering effect.

**Figure 3.**
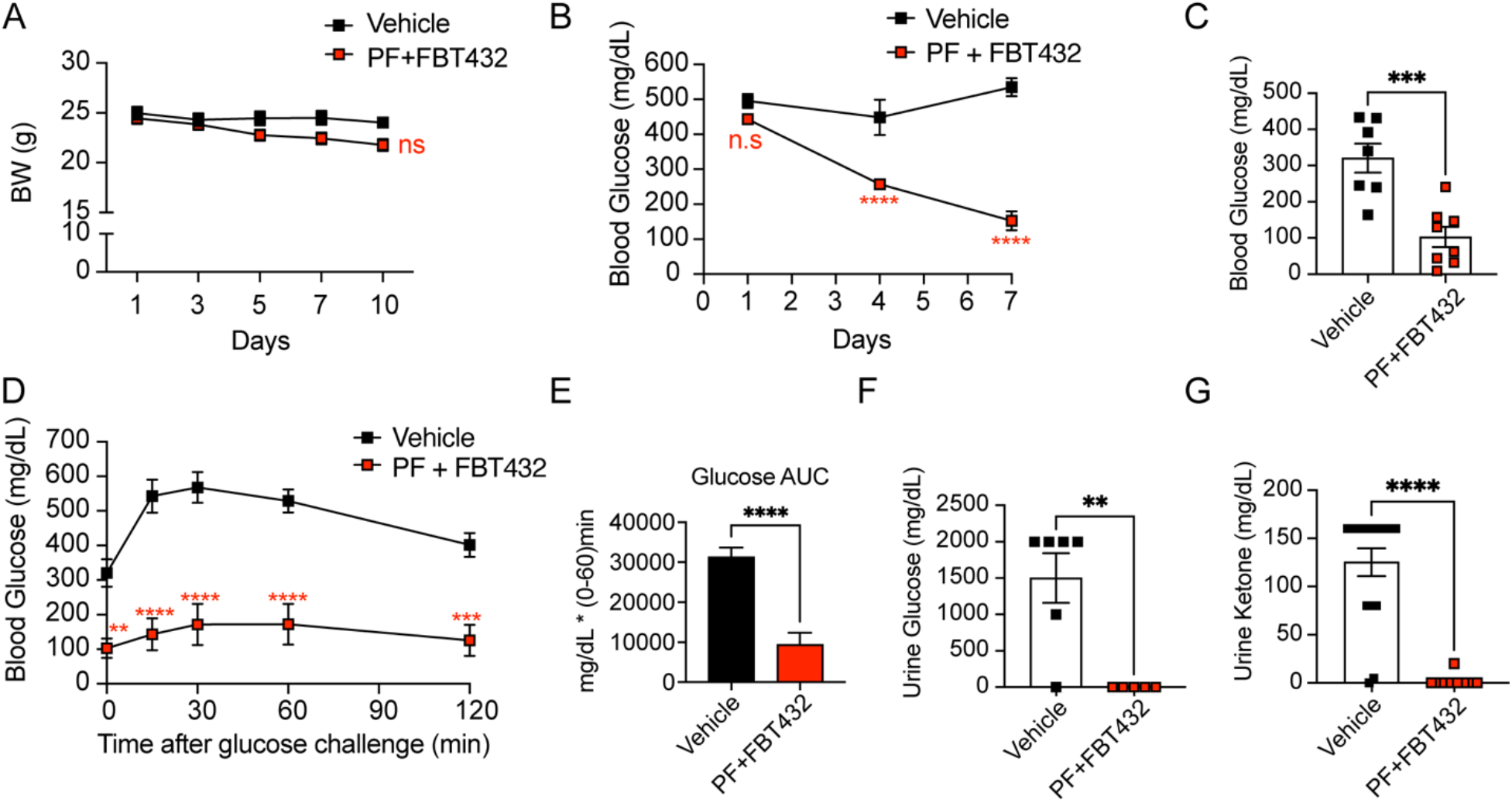
Combination treatment with PF-03084014 and FBT432 in STZ mice. (A) Weight measured before morning dosing. NS= not significant vs. vehicle by two-way ANOVA. N=8-10 per group. Data are means ± SEM. (B) *Ad libitum* glucose measured before afternoon dosing. ****= p < 0.0001, ns= not significant vs. vehicle by two-way ANOVA. (C) 4-hr fasting glucose on day 10. ***= p < 0.001 vs. vehicle by two-tailed t-test. (D) Glucose tolerance test (GTT) on day 10. *, ***, ****= p < 0.01, 0.001, 0.0001 vs. vehicle by two-way ANOVA. (E) AUC from 0-60 min during GTT. ****= p < 0.0001 vs. vehicle by two-tailed t-test. (F) Glycosuria and (G) ketonuria on day 10. **, ****= p < 0.01, 0.0001 vs. vehicle by two-tailed t-test.

**Figure 4.**
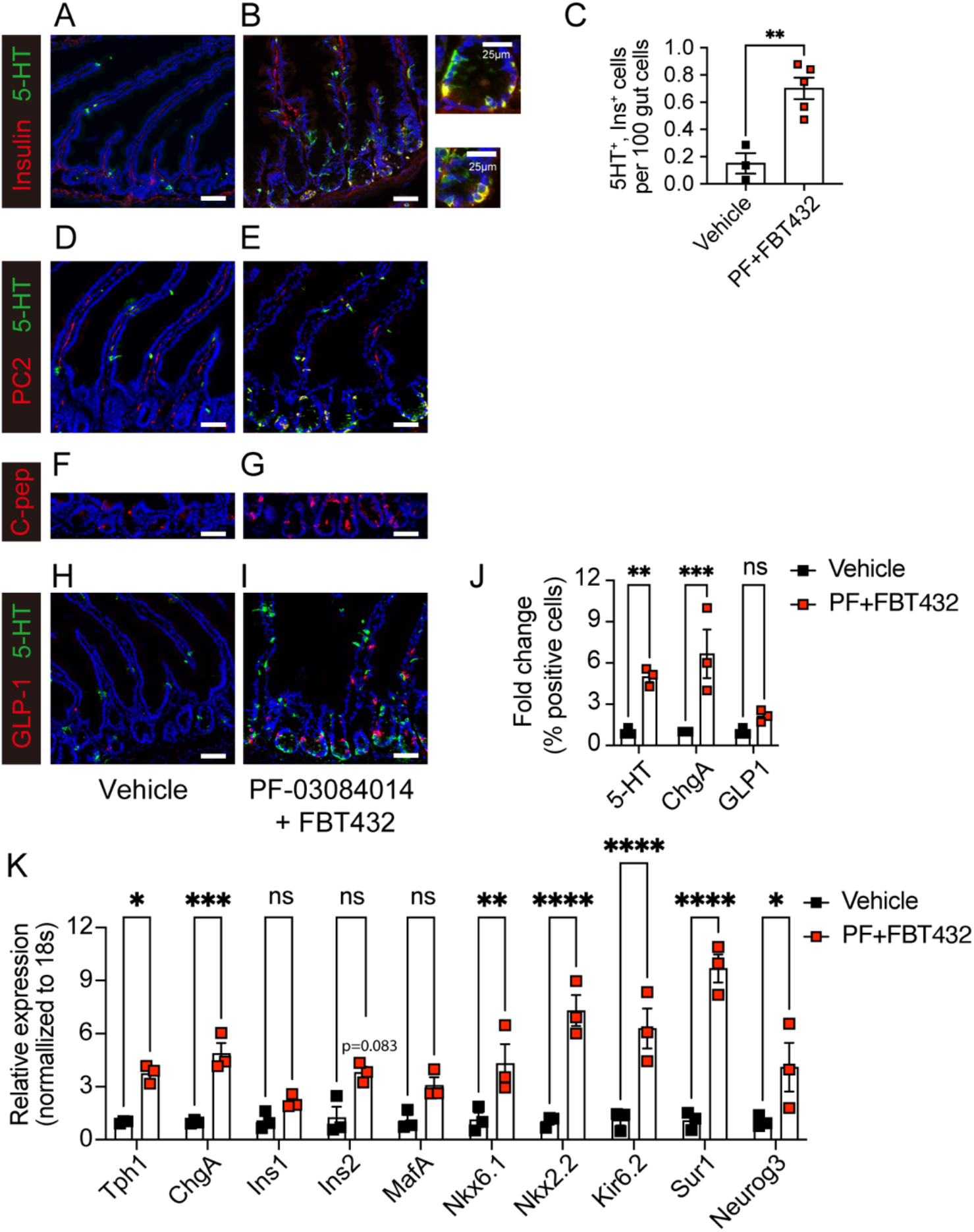
Combination treatment with PF-03084014 and FBT432 and β-like gut cells. (A, B) Representative intestinal immunostaining with 5HT (green) and insulin (red) after 10 days of treatment. (C) Quantification of 5HT- and insulin-immunoreactive cells across the entire gut. N=3-5 per group, data are means ±SEM. **= p < 0.01 vs. vehicle by two-tailed t-test. (D, E) 5-HT (green) and PC2 (red), (F, G) C-peptide, (H, I) 5-HT (green) and GLP-1 (red) staining. Scale bar= 50 μm, DAPI counterstains nuclei. (J) Flow cytometry of 5HT-, ChgA-, and GLP-1-immunoreactive cells from the proximal gut. N=3 per group, data are means ± SEM. **, ***= p < 0.01, 0.001 vs. vehicle by two-way ANOVA. (K) qPCR in isolated epithelial cells. N=3 per group, data are means ± SEM. *, **, ***, ****= p < 0.1, 0.01, 0.001, 0.0001 vs. vehicle by twoway ANOVA.

Lastly, we examined whether the glucose lowering effects of the combination treatment was due in part to insulin-mediated effects on hepatic glucose homeostasis. Gck expression was significantly increased in the liver during the combination treatment, while expression of gluconeogenic genes such as G6pc and Pck was suppressed compared to vehicle-treated group, consistent with a restoration of the actions of insulin, likely secreted from gut β-like cells (Figure 5A). Liver histology showed no lipid accumulation and normal glycogen stores (Figure 5B, C).

**Figure 5.**
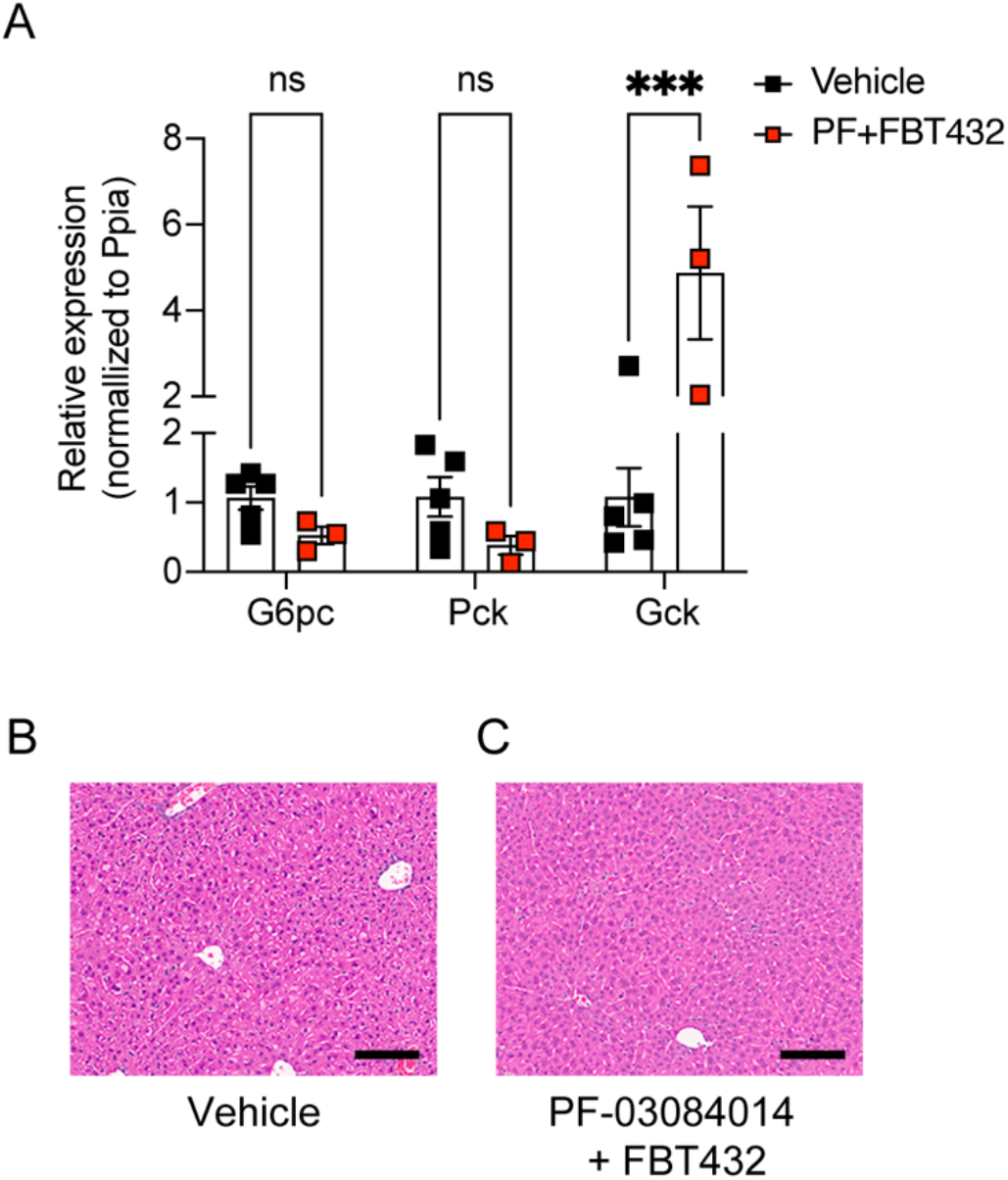
Hepatic effects of combination treatment with PF-03084014 and FBT432 in STZ mice. (A) RNA analysis after 10 days of treatment. (B) Representative H&E staining after 10 days of treatment. Scale bar= 50 μm. N=3 per group, data are means ± SEM. ***= p < 0.001, ns= not significant vs. vehicle by two-way ANOVA.

### Glucose lowering effect of FBT374 in STZ-induced diabetic mice

In addition to FBT432, we identified other active compounds with good oral bioavailability, suitable for progression to *in vivo* models, including FBT374. In activity and selectivity studies, FBT374 displayed a strong potency against FOXO1 (estimated IC_50_=73 nM), no activity against FOXA2, and minimal activity against FOXO3 and FOXO4. No cellular toxicity or nonspecific activity against CMV promoter-mediated reporter gene activity was observed (Table 2). We examined inhibition of FOXO1-dependent gluconeogenesis in primary hepatocytes by FBT374 and observed that it suppressed cAMP/DEX-induced *G6pc* in a dose-dependent manner with an estimated IC_50_ of 1.31 μM (Figure 6A).

**Figure 6.**
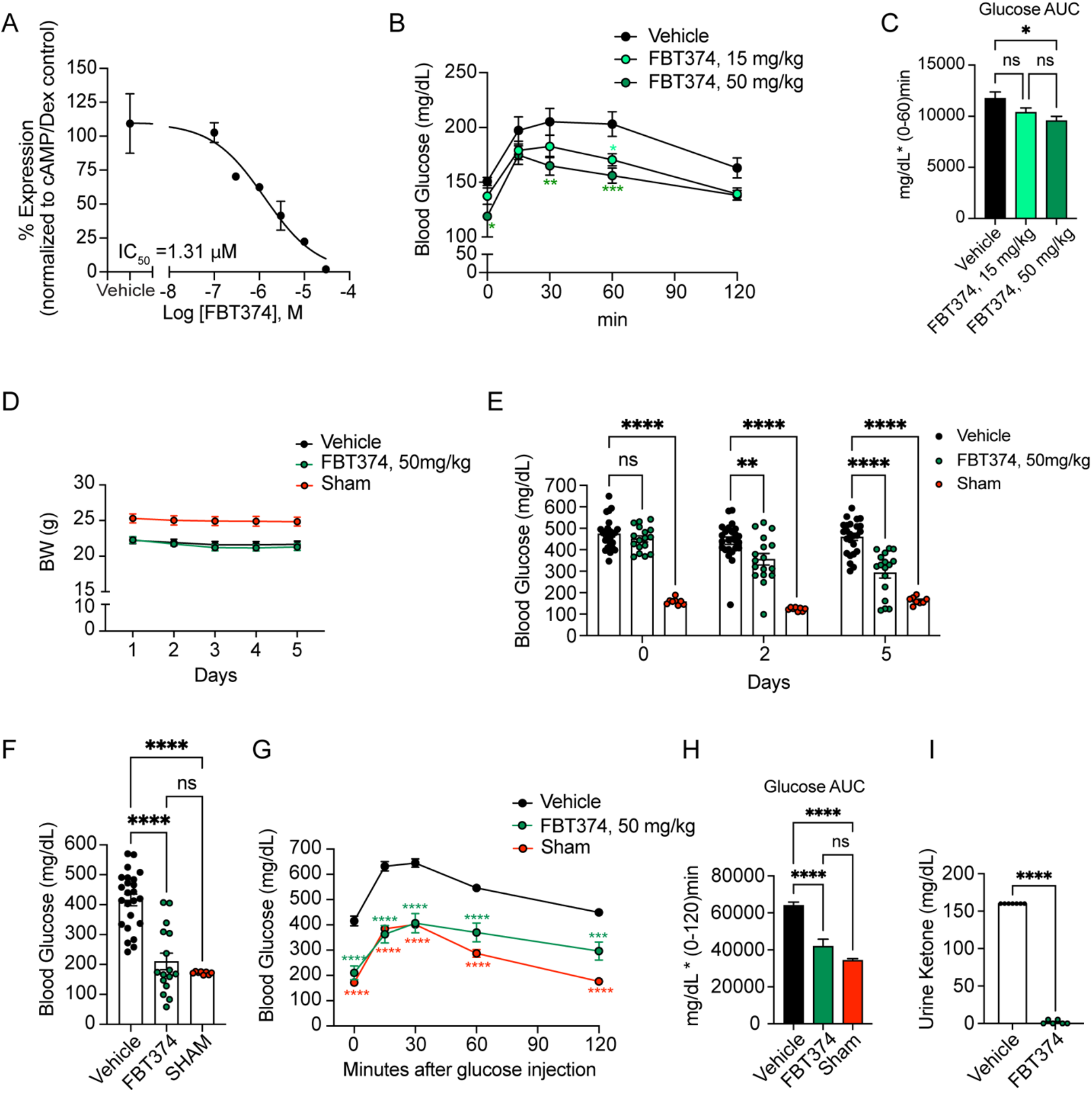
*In vitro* and *in vivo* activity of FBT374. (A) *G6pc* mRNA in hepatocytes treated with the combination described. Means ± SEM of 2-3 replicate wells per concentration are shown. IC_50_ was generated by four-parameter curve fit in GraphPad Prism. (B, C) Glucose levels and area-under-the-curve (AUC) during intraperitoneal PTT after oral treatment with FBT374. N=8 mice per group; *, **, ***= p < 0.05, 0.01, 0.001 vs. vehicle by two-way ANOVA (B) or one-way ANOVA (C). 7-8-week-old C57/BL6 mice were randomized into three groups based on the body weight. (D) Weight measured before morning dosing in STZ or control mice. NS= not significant by two-way ANOVA. N= 8-25 per group. Aggregated data from two independent experiments conducted under the same conditions and presented as means ± SEM. (E) *Ad libitum* glucose measured before afternoon dosing. **, ****= p < 0.01, 0.0001 vs. vehicle by two-way ANOVA. (F) 4-hr fasting glucose on day 6. ***= p < 0.001 by one-way ANOVA multiple comparison. (G) Glucose tolerance test (GTT) on day 6. ****= p < 0.0001 vs. vehicle by two-way ANOVA. (H) AUC from 0-120 min during GTT. **= p < 0.01 by one-way ANOVA for multiple comparisons. (I) Urine ketone on day 7. ****= p < 0.0001 vs. vehicle by two-tailed t-test.

*In vitro* DMPK studies showed that FBT374 is more cell permeable than FBT432 while retaining similar solubility (Supplementary Table 1). Despite lower *in vitro* metabolic stability than FBT432, FBT374 has comparable oral bioavailability. The slightly lower systemic plasma levels were accompanied by higher exposure in liver and small intestine, at both 1h and 8h after oral dosing at 50 mg/kg (supplementary Table 3). Together with the higher cell permeability, the exposure data prompted us to investigate whether FBT374 is effective as a single agent FOXO1 inhibitor.

To determine whether FBT374 suppresses hepatic glucose production (HGP) *in vivo*, we performed pyruvate tolerance tests (PTT) in normoglycemic mice following three oral doses of FBT374 at 15 and 50 mg/kg, respectively. Following intraperitoneal pyruvate injection, 50 mg/kg FBT374-treated animals showed a significantly lower glucose excursion than vehicle-treated controls; we also saw a similar trend in animals treated with 15 mg/kg (Figure 6B-C). Based on these results we further evaluated the glucose-lowering effect of FBT374 in STZ mice.

As a control we included a vehicle-treated group that underwent the same procedures as STZ-treated mice. Of note, STZ injection and uncontrolled hyperglycemia resulted in a ~10% weight loss, seen as early as day 1 (Figure 6D). Oral administration of FBT374 at 50 mg/kg/dose b.i.d. for 6 days was well tolerated without obvious signs of toxicity, consistent with the lack of cellular toxicity *in vitro*. Interestingly, a progressive and significant glucose lowering effect was observed within 2 days after starting treatment with FBT374 and persisted until the end of the study (Figure 6E). Nearly a third (5 out 17) of FBT374-treated mice achieved euglycemia compared to the vehicle group after 5 days of treatment, and 61% (10 out of 16) showed fasting euglycemia compared to sham-treated controls (Figure 6F). Strikingly, following intraperitoneal injection of glucose after a 4-hr fast, FBT374-treated mice exhibited a glucose excursion identical to the sham group at 15 and 30 min, and slightly higher levels at 60 and 120 min compared to sham (Figure 6G). Area under the curve (AUC) calculation confirmed that glucose tolerance was virtually normal following FBT374 treatment of STZ-diabetic mice (Figure 6H). Accordingly, ketonuria, an indicator of insulin-deficient diabetes, became undetectable (Figure 6I). These data suggest that FBT374 treatment alone can efficiently improve glucose homeostasis in STZ-induced diabetic mice.

To determine whether reprogramming of enteroendocrine cells into the β-like cells contributed to the improved glycemic control of FBT374-treated mice, we examined the presence of insulin-immunoreactive gut cells by co-immunostaining with insulin and 5HT. Insulin- and 5HT-immunoreactive cells were increased in duodenal crypts (Figure 7A, B). Quantification of the entire gut from 7 different FBT374-treated mice confirmed a ~5-fold increase of insulin-immunoreactive cells compared to controls (Figure 7C). This increase was paralleled by PC2- /5HT-, and C-peptide co-immunoreactive cells in FBT374-treated mice (Figure 7D-G). Consistent with our previous findings, gut immunohistochemistry showed that FBT374 treatment increased the 5HT-immunoreactive EEC subpopulation by ~2.5 fold, while GLP-1 cells remained unchanged compared to vehicle-treated mice (Figure 7J), indicating that there is an expansion of EECs capable of conversion to β-like cells (Figure 7H, I). Gene expression analysis confirmed a significant increase of *Tph1* with a trend to increase *ChgA* (Figure 7K).

**Figure 7.**
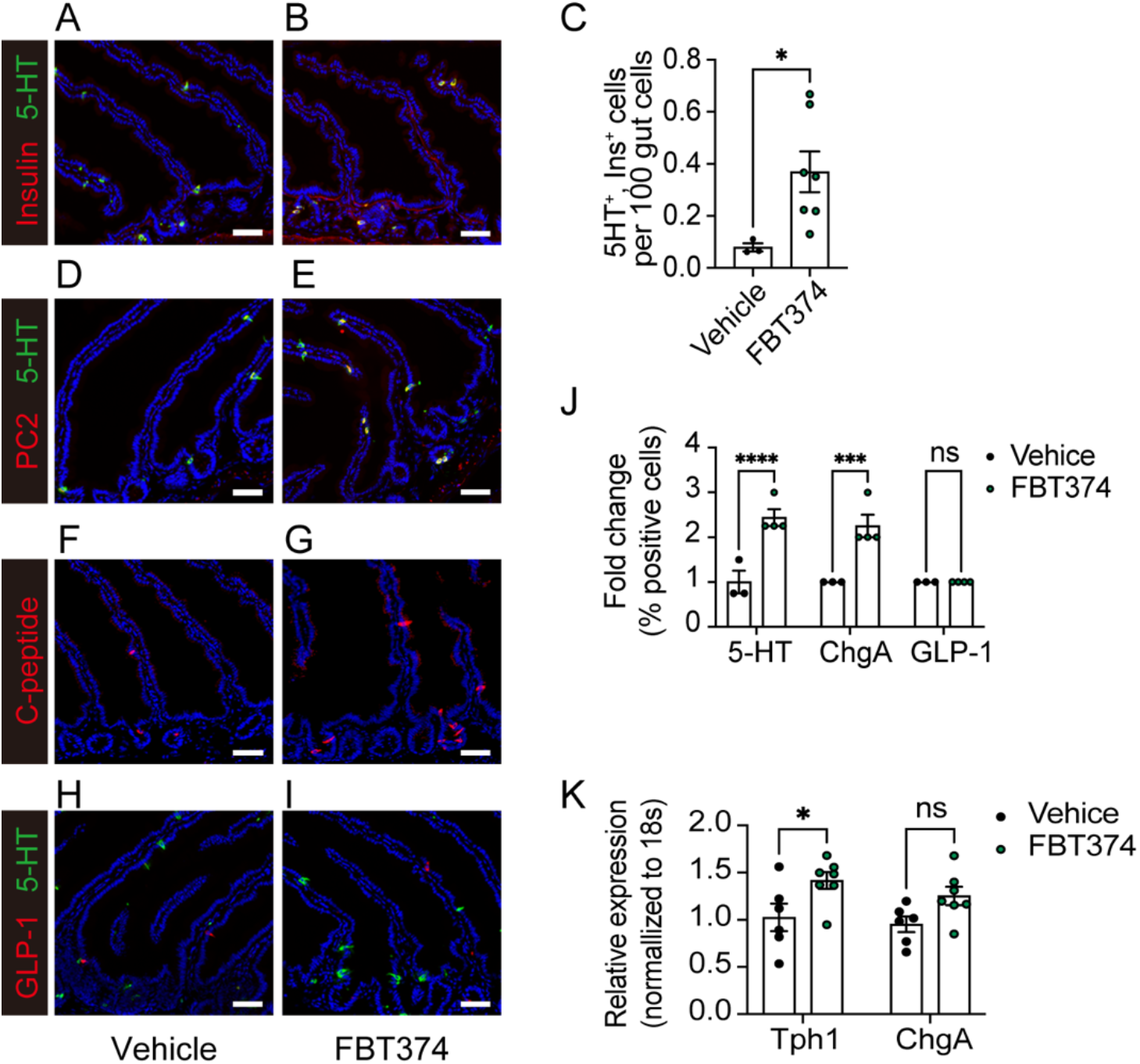
Combination treatment with FBT374 and generation of β-like cells. (A, B) Representative immunostaining with 5HT (green) and insulin (red) after 12-day treatment with vehicle or FBT374. (C) Quantification of 5HT- and insulin-immunoreactive cells across the entire gut. N=3-7 per group, data are means ± SEM. * p < 0.05 vs. vehicle by two-tailed t-test. (D, E) 5HT (green) and PC2 (red), (F, G) C-peptide, (H, I) 5HT (green) and GLP-1 (red) staining. Scale bar= 50 μm, DAPI counterstains nuclei. (J) Flow cytometry of 5HT-, ChgA-, and GLP-1-immunoreactive cells from the proximal gut. N=3-4 per group, data are means ± SEM. ***, ****= p < 0.001, 0.0001 vs. vehicle by two-way ANOVA. (K) qPCR of isolated epithelial cells. N=6-7 per group, data are means ± SEM. * =p < 0.1, **= p< 0.01, ns= not significant vs. vehicle by two-way ANOVA.

## Discussion

We report the identification and characterization of potent, selective, and orally bioavailable FOXO1 inhibitors capable of inducing conversion of gut cells to insulin-producing, glucose-responsive β-like cells capable of normalizing glucose homeostasis in insulin-deficient diabetic mice. We optimized a series of small molecule FOXO1 inhibitors, confirming their activity and selectivity in reporter gene and primary hepatocyte functional assays. We advanced compounds with favorable *in vitro* ADME properties and *in vivo* PK, including FBT432 and FBT374. When administered to mice, both compounds decreased glucose excursions during pyruvate tolerance tests, consistent with the reported effects of FOXO1 on glycogenolysis and gluconeogenesis (15). This decrease was accompanied by suppression of cAMP/dexamethasone-induced *G6pc* expression in primary hepatocytes, constituting effective, combined pharmacodynamic readouts of compound efficacy.

The effects of treatment with FBT432 in combination with PF-03084014 are reminiscent of those in Akita mice by a combination of FOXO1 and Notch inhibitors (10). These findings suggest that the ability to expand the EEC pool and thus the generation of cells with an insulinogenic potential is a class effect of this combination. Given the demonstrated *in vivo* safety of various Notch inhibitors in human trials for indications from Alzheimer’s disease to cancer, and the obvious benefits of an oral treatment that would liberate patients from devices to release insulin and measure glucose–let alone the constant risk of fatal hypoglycemia–this approach has merit.

However, the main advance of the present study is the demonstration of efficacy of FOXO1 inhibition by a single compound. In this regard, FBT374 outperformed FBT432. With both FBT compounds, we found a correlation between hypoglycemic effect and number of insulin-immunoreactive gut cells. While comparable to FTB432 in primary luciferase assays, FBT374 showed improved solubility and cellular permeability and, despite lower *in vitro* metabolic stability, good oral bioavailability, tissue exposure, and higher intestinal levels than FBT432, which may lead to better local exposure in the crypts, where we observed the most pronounced effect on the generation of β-like cells. FBT374 as monotherapy normalized glucose levels and glucose tolerance in STZ diabetes, a finding that recapitulates both somatic ablation of Foxo1 by Neurogenin3 *Cre* and combination treatment.

These results phenocopy those obtained in STZ mice following genetic ablation of Foxo1 and are consistent with the possibility that gut insulin production drives the therapeutic effect (8). While we cannot rule out a contribution to the glucose-lowering effect by reduced hepatic glycogenolysis and gluconeogenesis, this effect would be therapeutically desirable and by itself cannot account for the normalization of glycemia, especially during the early time points following glucose administration, when insulin-induced glucose uptake in muscle and adipose predominates over suppression of hepatic glucose production (16–18). In addition, the reversal of ketonuria can only be explained by inhibition of lipolysis, another effect of insulin that cannot be ascribed to improved insulin sensitivity. We were unable to detect an increase of plasma insulin, likely for two reasons: unlike in the pancreas, where β-cells are clustered together and release insulin in a coordinated fashion, β-like cells in the gut are anatomically scattered, resulting in a shallower secretory peak. Moreover, given the insulin-deficient state, secreted insulin is likely to be rapidly cleared by the liver, resulting in lower systemic concentrations (19–21).

The feasibility of converting gut cells into β-like cells has received further support by the recent discovery of *bona fide* insulin-producing cells in the human fetal gut (22), an observation that we have independently confirmed (23). In this regard, it is possible that FOXO1 inhibition arrests EEC maturation at a fetal stage of development, a speculation that is entirely consistent with the known effects of FOXO1 inhibition in the pancreatic islet, where it promotes reversion to a progenitor-like stage (24; 25). Thus, this intervention may recapitulate aspects of the normal ontogeny of the cell fates in the endodermal anlagen from which intestine, liver and islets are derived.

In conclusion we have identified potent, selective, orally bioavailable Foxo1 inhibitors that show efficacy in normalizing glucose levels in STZ diabetes, either in combination with a Notch inhibitor (thereby extending our previous findings in the Akita model) (10) or as single agents, phenocopying the effect of genetic Foxo1 ablation in mouse (8). Our results suggest the use of Foxo1 inhibitors as a potential new class of agents for the treatment of insulin-dependent diabetes.

## Acknowledgements

We thank Drs. Utpal B. Pajvani and Joel Berger for insightful discussions of the data, and Ms. Ana Maria Flete, and the Graduate Center Advanced Science Research Center of the City University of New York for exceptional technical support.

## Author contribution

Y.L designed study, developed methodology, acquired data, analyzed data, and wrote manuscript. Y.N performed experiment, analyzed data, and wrote manuscript. B.D and N.S performed experiment and contributed to discussion. T.K and W.D contributed to discussion. R.L and A.D reviewed/edited the manuscript. S.B supervised the study, wrote manuscript.

## Funding and others

This work was partly funded by a sponsored research agreement between Forkhead BioTherapeutics, Inc. and Columbia University (RL), by SBIR grant R44DK120177, and by grants P30CA013696 and P30DK63608 through the resources of the Flow Core Facility.

---

**Supplementary Figure 1.**
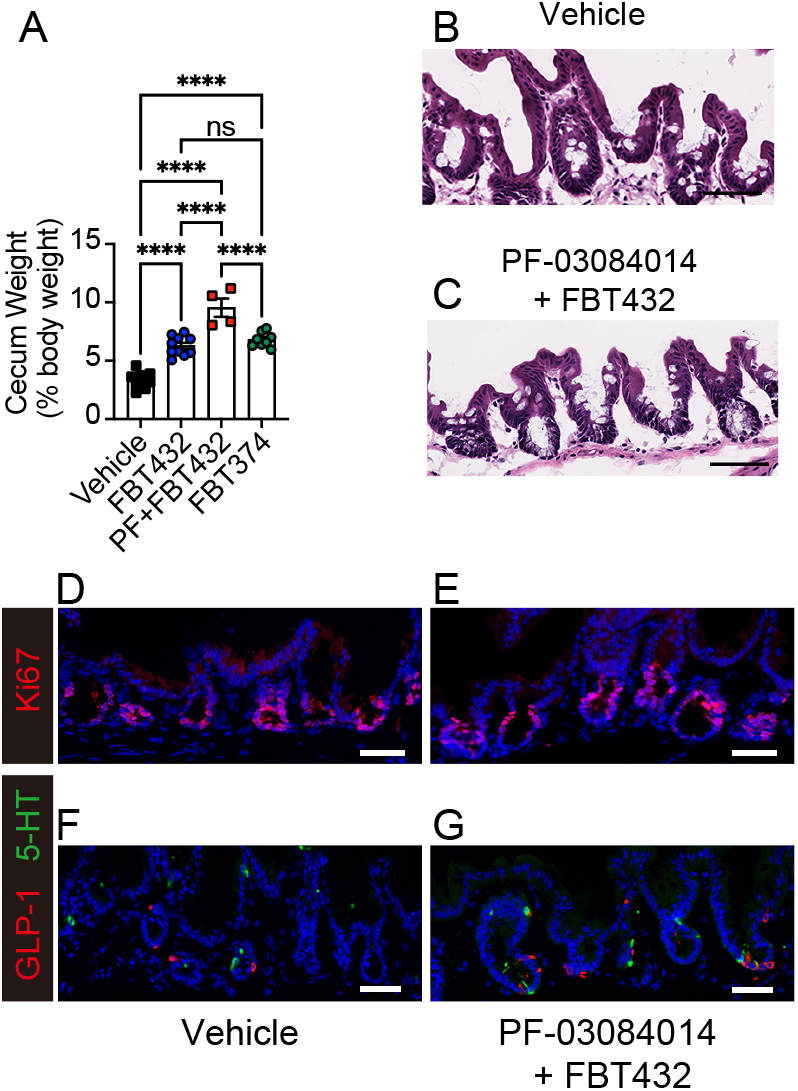
Enlarged cecum in FBT374 or FBT432 or PF-0308014+FBT432 treated mice. (A) A wet cecum weight normalized by body weight. N = 4-10 per group. ****= p < 0.0001; ns= not significant by one-way ANOVA multiple comparison. (B) Representative images of H&E or immunostaining of cecum collected from vehicle- or PF-030804014+FBT432-treated mice. (B, C) Sections were stained with hematoxylin and eosin, (D, E) Ki67, (F, G) 5HT and GLP-1 antibodies. Images were taken using the Keyence BZ-X800 microscope. Scale bar= 50 μm, N=3 per group.

## Supplementary tables

**Supplementary Table 1.** *In vitro* DMPK.

|  |  | FBT432 | FBT10 | FBT374 |
| --- | --- | --- | --- | --- |
| Mouse microsomal stability | Clint: (ul/min/mg protein) | < 10 | 165 | 195.8 |
| Cell Permeability MDCKII | Papp average (1e-6 cm/s) | 4.86 | 5.9 | 10.3 |
|  | B-A/A-B ratio | 0.8 | 1.4 | 1.6 |
| Solubility | PBS. pH7.4 mean (mM) | 70.5 | 5.3 | 80.3 |

**Supplementary table 2.** FBT-432 *in vivo* PK and tissue distribution.

| **FBT-432 in plasma after IV and PO at 1 mg/kg and 10 mg/kg in male ICR mice** |  |  |
| --- | --- | --- |
| <u>IV</u>, Single administration, 1 mg/kg |  |  |
| Parameters | Unit | Mean |
| t<sub>1/2</sub> | h | 1.48 |
| AUC 0-inf obs | mM*h | 1.02 |
| Cl obs | ml/min/kg | 38.4 |
| V<sub>ss</sub> obs | L/kg | 3.22 |
| <u>PO</u>, Single administration, 10 mg/kg |  |  |
| Parameters | Unit | Mean |
| t<sub>1/2</sub> | h | 2.01 |
| T<sub>max</sub> | h | 0.5 |
| C<sub>max</sub> | mM | 0.79 |
| AUC 0-inf obs | mM*h | 3.41 |
| *F | % | 33.5 |

| **FBT-432 in plasma after PO administration at 50 mg/kg in male ICR mice** |  |  |
| --- | --- | --- |
| PO, Single administration, 50 mg/kg |  |  |
| Parameters | Unit | Mean |
| t<sub>1/2</sub> | h | 2.78 |
| T<sub>max</sub> | h | 2.67 |
| C<sub>max</sub> | mM | 3.90 |
| AUC 0-inf_obs | mM*h | 28.11 |
| *F | % | 55.3 |

**Tissue distribution parameters of FBT-432 after PO administration at 50 mg/kg in male ICR mice**
| Sampling Time (h) | Matrix | Mean Conc. | Tissue / Plasma ratio |
| --- | --- | --- | --- |
|  |  | (mM) |  |
| 1 | Plasma | 2.11 | - |
|  | Brain | 0.06 | 0.03 |
|  | Liver | 29.27 | 13.86 |
|  | Kidney | 18.21 | 8.62 |
|  | Small intestine | 17.49 | 8.28 |
|  | Large intestine | 445.98 | 211.22 |
| 8 | Plasma | 1.19 | - |
|  | Brain | 0.13 | 0.11 |
|  | Liver | 15.01 | 12.62 |
|  | Kidney | 11.57 | 9.73 |
|  | Small intestine | 20.81 | 17.50 |
|  | Large intestine | 487.79 | 410.16 |

**Supplementary table 3.**
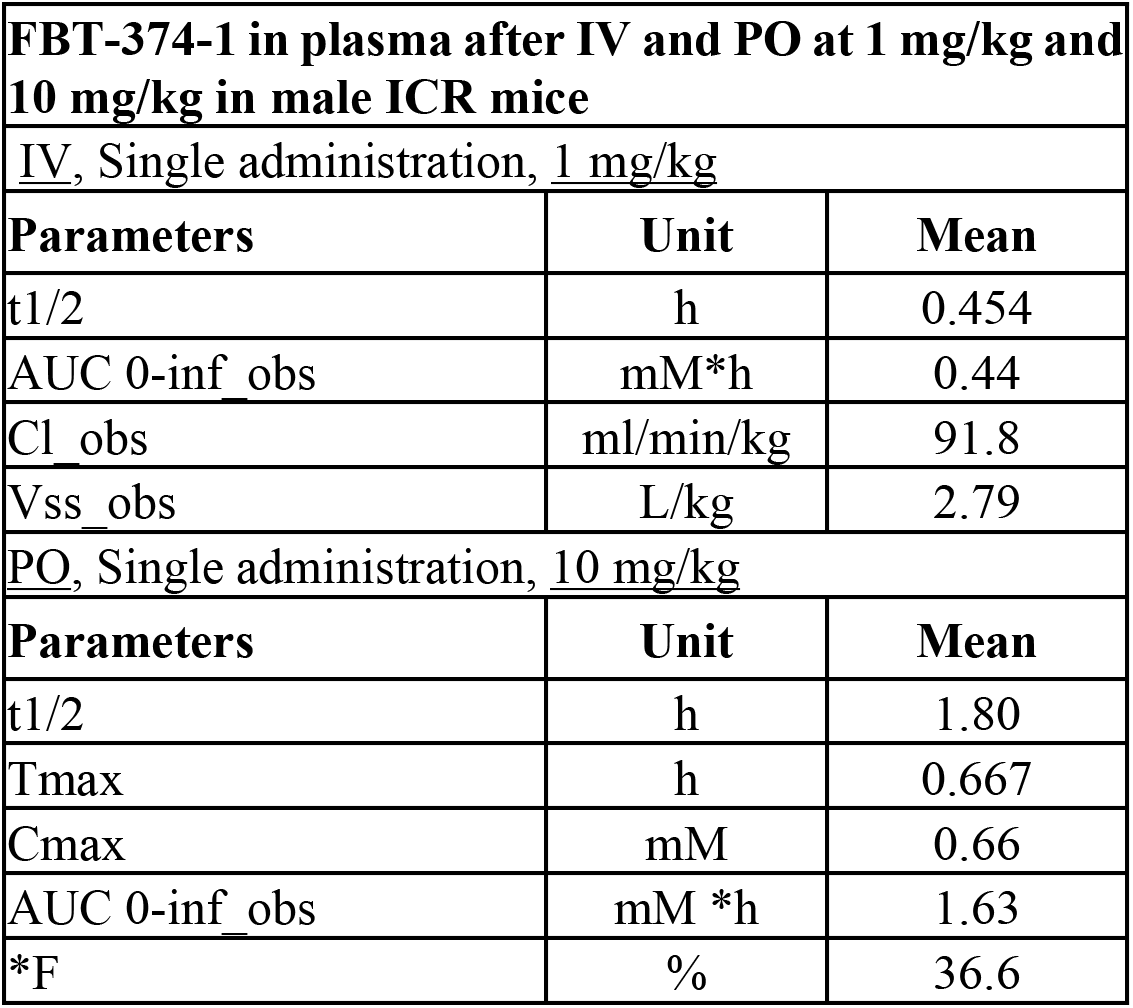
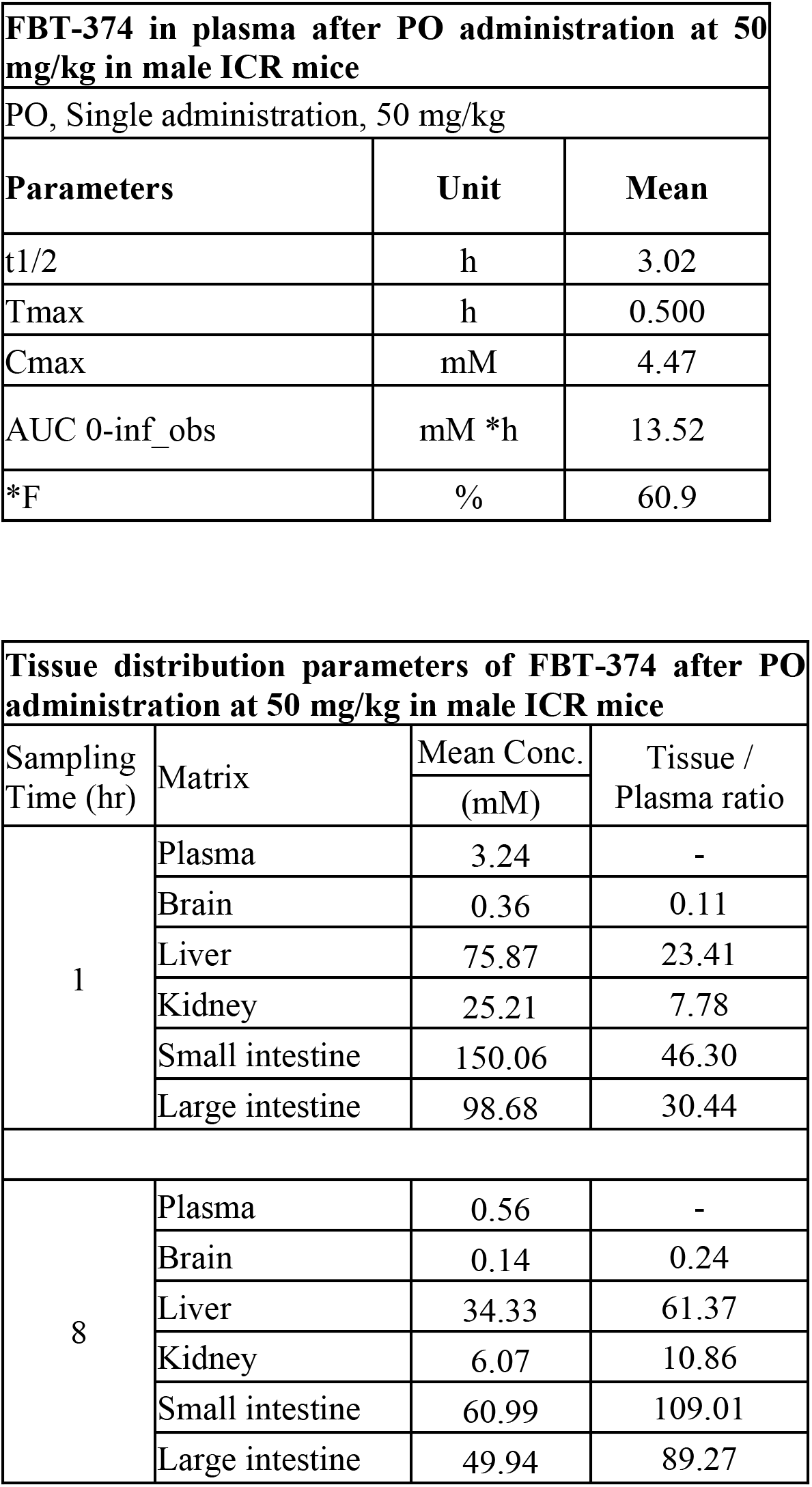
FBT-374 *in vivo* PK and tissue distribution.

**Supplementary table4.**
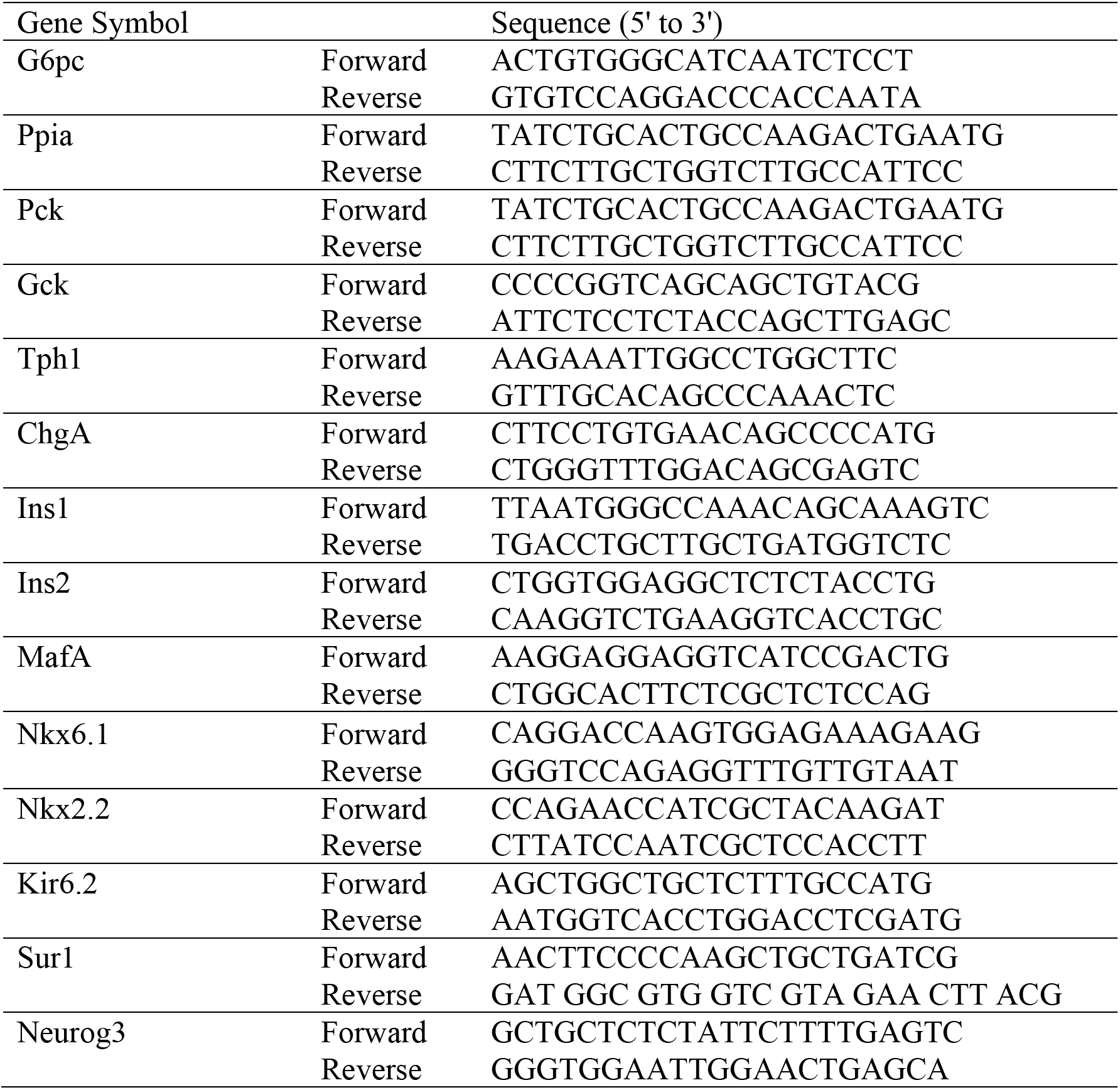
List of primers for qPCR.

**Supplementary table5.**
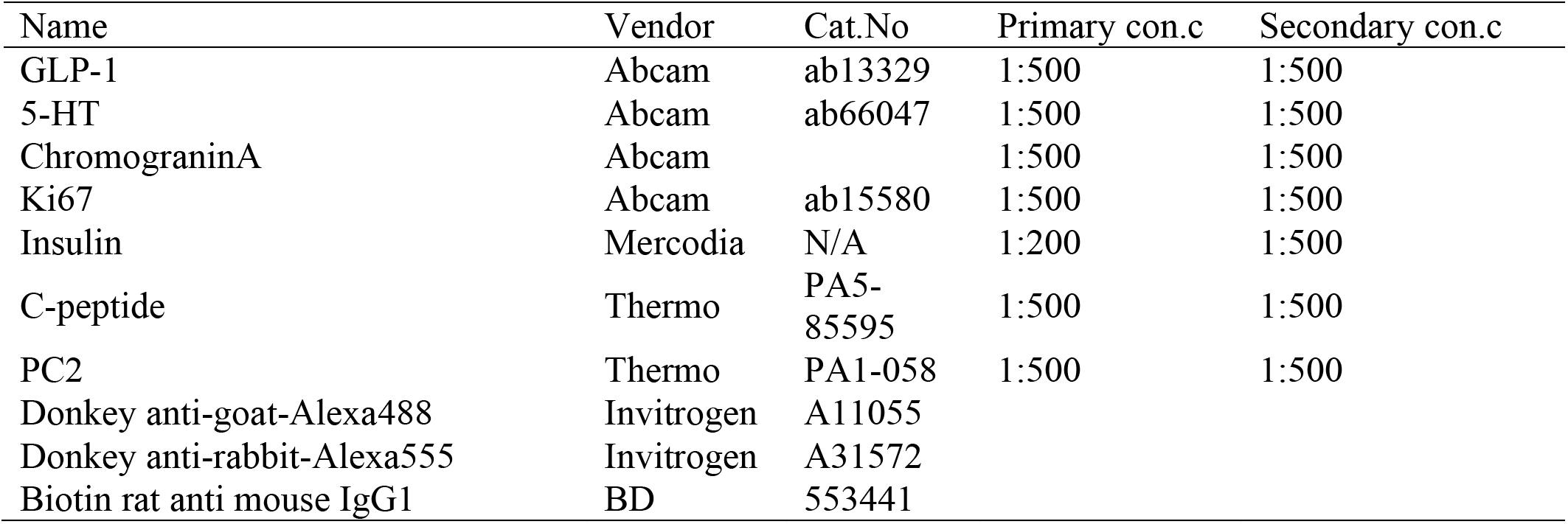
List of antibodies for immunohistochemistry or intracellular staining for flow cytometry analysis.

